# Selective role of the translin/trax RNase complex in hippocampal synaptic plasticity

**DOI:** 10.1101/2020.05.19.105189

**Authors:** Alan Jung Park, Mahesh Shivarama Shetty, Jay M. Baraban, Ted Abel

**Affiliations:** Department of Biology, University of Pennsylvania, Philadelphia, Pennsylvania; Department of Neuroscience and Pharmacology, Carver College of Medicine, University of Iowa, Iowa City, Iowa, United States; Iowa Neuroscience Institute, Carver College of Medicine, University of Iowa, Iowa City, Iowa, United States; The Solomon H. Snyder Department of Neuroscience, Johns Hopkins University School of Medicine, Baltimore, Maryland, United States; Mortimer B. Zuckerman Mind Brain Behavior Institute, Columbia University, New York, United States

**Author notes:** corresponding authors **Addresses for Correspondence:** Ted Abel, Iowa Neuroscience Institute, Department of Neuroscience and Pharmacology, Carver College of Medicine, University of Iowa, 2312 Pappajohn Biomedical Discovery Building, 169 Newton Road, Iowa City, Iowa 52242-1903, USA., Alan Jung Park, Gogos Lab, Columbia University, Mortimer B. Zuckerman Mind Brain Behavior Institute, Jerome L. Greene Science Center, L5-053, 3227 Broadway, New York, NY 10027, USA.

**Keywords:** Translin, trax, long-term potentiation, long-term depression, local protein synthesis, hippocampal synaptic plasticity, FMRP, RNA-binding protein, microRNA

## Abstract

Activity-dependent local protein synthesis is critical for synapse-specific, persistent plasticity. Abnormalities in local protein synthesis have been implicated in psychiatric disorders. We have recently identified the translin/trax microRNA-degrading enzyme as a novel mediator of protein synthesis at activated synapses. Additionally, mice lacking translin/trax exhibit some of the behavioral abnormalities found in a mouse model of fragile X syndrome. Therefore, identifying signaling pathways interacting with translin/trax to support persistent synaptic plasticity is a translationally relevant goal. Here, as a first step to achieve this goal, we have assessed the requirement of translin/trax for multiple hippocampal synaptic plasticity paradigms that rely on distinct molecular mechanisms. We found that mice lacking translin/trax exhibited selective impairment in a form of persistent hippocampal plasticity, which requires postsynaptic PKA activity. In contrast, enduring forms of plasticity that are dependent on presynaptic PKA were unaffected. Furthermore, these mice did not display exaggerated metabotropic glutamate receptor-mediated long-term synaptic depression, a hallmark of the mouse model of fragile X syndrome. Taken together, these findings demonstrate that translin/trax mediates long-term synaptic plasticity that is dependent on postsynaptic PKA signaling.

## Introduction

Extensive evidence suggests that localization of key mRNAs in the vicinity of synapses and activity-mediated regulation of their translation contributes to persistent forms of synaptic plasticity related to long-term memory (1-4). This synapse-specific, activity-dependent mechanism requires precisely regulated molecular signaling in presynaptic or postsynaptic compartments (2, 3, 5). Several RNA-binding proteins (RBPs) take part in the trafficking and translational regulation of specific mRNAs and thus allowing diversity in the mechanisms engaged by different forms of plasticity (6-8). The RNA-binding protein translin is an evolutionarily conserved brain-enriched protein, which regulates RNA trafficking and translational control (9-12). Together with its partner protein, translin-associated factor X (trax), these proteins form a microRNA-degrading enzyme that can trigger protein synthesis by reversing microRNA-mediated silencing (13-15). We have previously shown that the translin/trax RNase complex mediates activity-dependent local synaptic protein synthesis required for input-specific heterosynaptic plasticity (synaptic tagging and capture) and memory formation (15). However, the mechanisms that regulate translin/trax activity within synaptic compartments have not been investigated.

Our previous findings suggest that translin/trax may interact with the cAMP-PKA signaling pathway. Specifically, the microRNA targets of translin/trax are predicted to regulate the expression of PKA-anchoring proteins, cAMP-degrading phosphodiesterases (PDEs), cAMP-producing Gs-coupled β2-adrenergic receptors and adenylyl cyclases (15). In fact, the cAMP-PKA signaling pathway is highly localized within presynaptic or postsynaptic compartments by PKA-anchoring proteins and mediates protein synthesis required for persistent synaptic plasticity and memory formation (16-20). Therefore, investigating whether translin/trax interacts with PKA signaling within presynaptic or postsynaptic compartments can provide clues to understanding molecular mechanisms linking translin/trax to synaptic plasticity and memory formation.

In the present study, we determined the role of translin/trax in distinct forms of synaptic plasticity, which require presynaptic or postsynaptic PKA signaling. As trax protein is unstable in the absence of translin, in the current study we used translin KO mice, which lack both translin and trax proteins (15, 21).

## Materials and Methods

All experiments were performed according to the National Institutes of Health guidelines and were fully approved by the Institutional Animal Care and Use Committee at the University of Pennsylvania.

### Translin knockout (KO) mice

The generation and maintenance of translin KO mice (MGI:2677496) were described previously (21, 22). Mice were backcrossed to C57BL/6J for more than 15 generations. Heterozygous male and heterozygous female mice were mated to produce homozygous translin KO mice and wildtype littermates. Mice were maintained on a 12h light/12h dark cycle with lights on at 8 am (ZT0). Food and water were available *ad libitum*. All experiments were performed during the light cycle using translin KO mice and wildtype littermates as controls. 2-to 5-month-old mice were used for all experiments except for the LTD experiments in which 4-to 6-week-old mice were used.

### Drugs

Forskolin (FSK, Molecular grade, Sigma), an adenylyl cyclase activator, was freshly prepared as a 50 mM solution in 100% ethanol and delivered at 50 μM final concentration in artificial cerebrospinal fluid (aCSF) as described before {Park, 2014 #22}. (*RS*)-3,5-Dihydroxyphenylglycine (DHPG, Tocris), a potent agonist of group I metabotropic glutamate receptors (mGluRs), was freshly prepared as a 10 mM solution in milliQ water and delivered at 100 μM final concentration in aCSF as previously described (23).

### Electrophysiology

Experiments were performed as described (15). Briefly, both male and female 2-5-month-old mice were sacrificed by cervical dislocation and hippocampi were quickly collected in chilled, oxygenated aCSF (124 mM NaCl, 4.4 mM KCl, 1.3 mM MgSO_4_⋅7H_2_O, 1 mM NaH_2_PO_4_⋅H_2_O, 26.2 mM NaHCO_3_, 2.5 mM CaCl_2_⋅2H_2_O and 10 mM D-glucose) bubbled with 95% O_2_ / 5% CO_2_. 400 μm-thick transverse hippocampal slices were prepared using a manual slicer (Stoelting) and placed in an interface recording chamber at 28ºC (Fine Science Tools, Foster City, CA). The slices were constantly perfused with aCSF at 1 ml/min (or 2.5 ml/min for the mGluR-LTD experiment). Slices were equilibrated for at least 2 hours in aCSF. The stimulus intensity was set to elicit ∼40% of the maximum field-EPSP amplitude determined by an input-output curve in each experiment. The first 20-min baseline values were averaged, and the average was used to normalize each initial fEPSP slope. The input–output relationship and paired-pulse facilitation (PPF) were measured as previously described {Park, 2017 #46}. To electrically induce long-term potentiation (LTP), spaced 4-train (four 1 s 100 Hz trains delivered 5 minutes apart), massed 4-train (four 1 s 100 Hz trains delivered 5 seconds apart), theta-burst stimulation (TBS, 15 bursts of four 100 Hz pulses delivered for a total of 3 s at 5 Hz), and one-train (one 1 s 100 Hz train) stimulation were delivered after 20 min baseline recordings. To chemically induce LTP, 50 μM of FSK in aCSF was bath applied to the slices for 15 minutes following 20-min baseline recordings. To chemically induce LTD, 100 μM of DHPG in aCSF was bath applied to the slices for 10 minutes following 20-min baseline recordings.

### Western blotting

Hippocampal tissue homogenization, protein separation and transfer to polyvinylidene difluoride (PVDF) membranes were performed as previously described (24). Membranes were blocked in 5% BSA or 5% non-fat milk in TBST and incubated with primary antibodies (translin,1:100,000; FMRP, 1:10,000, Millipore) overnight at 4°C. They were washed and incubated with appropriate horseradish peroxidase-conjugated goat anti-mouse or anti-rabbit IgG (1:10,000, Santa Cruz) for 1 h in room temperature. Blots were exposed on a film by ECL and quantified using ImageJ. The density of signal was normalized to beta-tubulin levels (1:50,000, Sigma). Translin antibody was produced (New England Peptide, Inc.) based on the sequences provided previously (25). The antibody synthesis was based on the C-terminal sequence of the human translin (CKYDLSIRGFNKETA).

### Data Analysis

Data analyses were performed using Statistica 10. The maintenance of LTP or LTD was analyzed using a two-way repeated-measures ANOVA test on the last 20-min of the recordings {Park, 2014 #22;Park, 2017 #46}. The average of the normalized slopes over the last 20-min was compared between two groups using unpaired t-test. Western blotting data was analyzed using unpaired t-test. The ‘n’ used in all the experiments represents the number of mice. Differences were considered statistically significant when *p* < 0.05. Data are plotted as mean ± S.E.M.

## Results

### Translin KO mice show deficits in a specific form of PKA-dependent long-lasting LTP

In our previous study, we found that translin knockout mice display normal basal synaptic transmission measured by paired-pulse facilitation and input-output curves. These mice also exhibit unaltered transient potentiation, induced by a single 100 Hz stimulation, that requires neither PKA activity nor protein synthesis (15). In the present study, we first tested long-lasting forms of LTP induced by spaced 4-train (four 100 Hz trains of 1 s each, delivered 5 minutes apart) or massed 4-train (four 100 Hz trains of 1 s each, delivered 5 s apart) stimulation. The latter does not depend on PKA activation, whereas the former requires postsynaptic PKA activity (26-30). Hippocampal slices from translin KO mice showed marked impairment in spaced 4-train LTP (**Fig. 1*B***; n = 5 for each group, two-way repeated-measures ANOVA, F_(1,8)_ = 34.43, *p* = 0.00038). The average of the initial fEPSP slope over the last 20 min of the recordings was reduced in slices from translin KO mice compared to slices from wildtype littermates (wildtype littermates: 176.6 ± 13.5%, n = 5; translin KO mice: 89.5 ± 9.6%, n = 5, t-test, *p* = 0.00037). On the other hand, massed 4-train LTP was unaltered in slices from translin KO mice as shown in **Fig. 1*C*** (n = 5 for translin KO mice, n = 5 for wildtype littermates, two-way repeated-measures ANOVA, F_(1,8)_ = 0.923, *p* = 0.365). The average of the initial fEPSP slope over the last 20 min of the recordings was similar between slices from translin KO mice and wildtype littermates (wildtype littermates: 143.6 ± 8.2%, n = 5; translin KO mice: 154.6 ± 9.9%, n = 5, t-test, *p* = 0.364).

**Figure 1.**
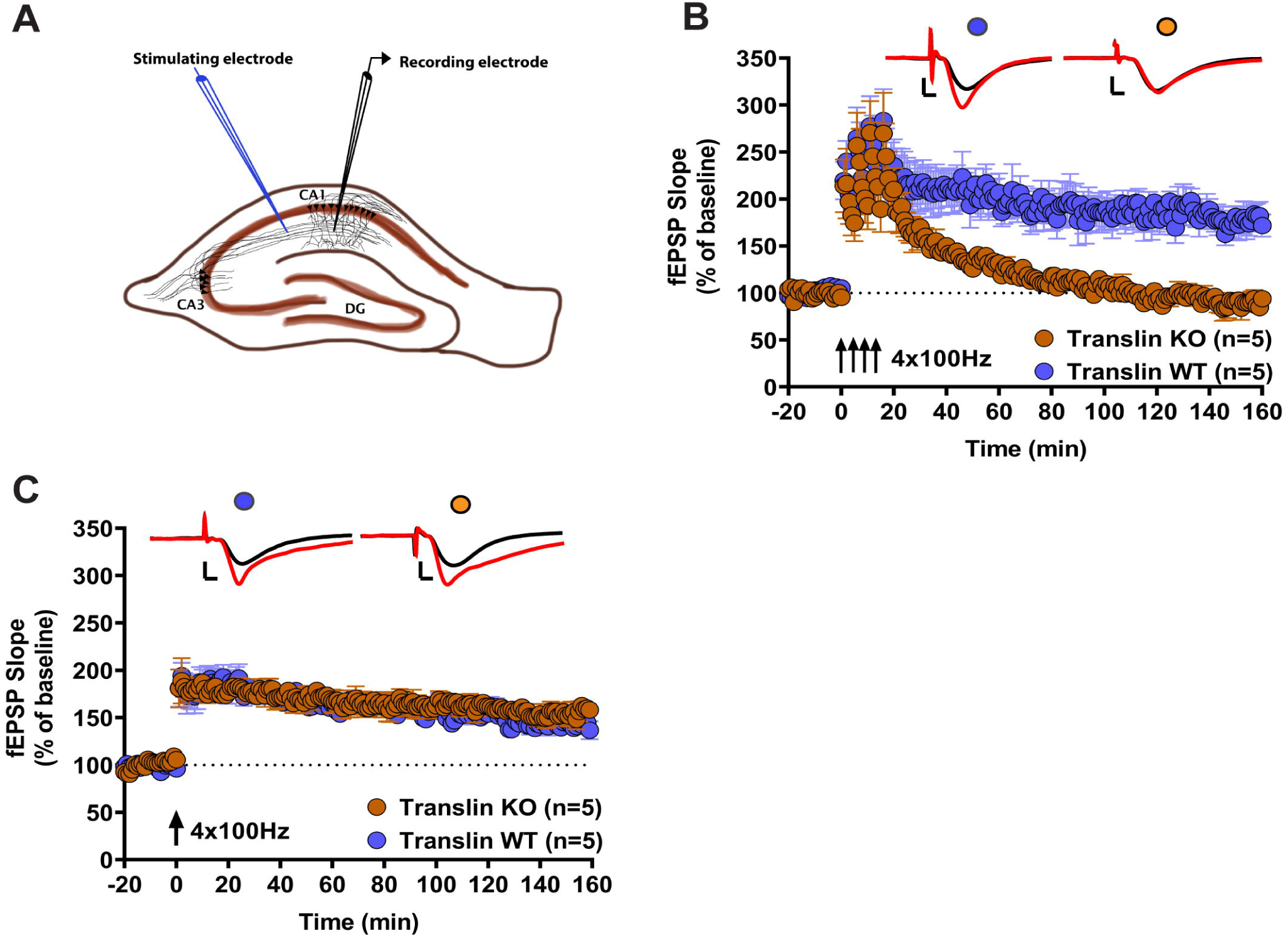
Translin KO mice exhibit deficits in a postsynaptic PKA-dependent form of long-lasting LTP. ***A***. A schematic representation of a hippocampal slice showing the placement of electrodes for field-EPSP recordings in the CA1 stratum radiatum upon stimulation of Schaffer collaterals. ***B***. Hippocampal slices from translin KO mice (n=5; 3 males, 2 females) showed impaired long-lasting LTP induced by spaced 4-train stimulation (four 1s 100 Hz stimuli delivered 5 minutes apart) compared to slices from wildtype littermates (n=5; 1 male, 4 females) (two-way repeated-measures ANOVA, F_(1,8)_ = 34.43, *p* = 0.00038). ***C***. Massed 4-train stimulation (four 1 s 100 Hz trains delivered 5 seconds apart) elicited long-lasting LTP that was not significantly different between slices from translin KO mice (n=5; 2 males, 3 females) and wildtype littermates (n=5; 1 male, 4 females) (two-way repeated-measures ANOVA, F_(1,8)_ = 0.923, *p* = 0.365). Representative traces before (black) and after (red) stimulation are shown on top of each graph. Scale bars for traces 2 mV, 2 ms. ‘n’ refers to the number of mice used. Error bars reflect S.E.M.

Next, we examined two other long-lasting forms of LTP induced by either theta-burst stimulation (TBS; 15 bursts of four 100 Hz pulses delivered at 5 Hz) or bath application of the adenylyl cyclase activator forskolin (FSK). These forms of LTP rely on increased transmitter release and require presynaptically compartmentalized PKA signaling (19, 28, 31-33). TBS-LTP was unaffected in slices from translin KO mice (**Fig. 2*A***; n = 5 for each group, two-way repeated-measures ANOVA, F_(1,8)_ = 0.007, *p* = 0.94). The average of the initial fEPSP slope over the last 20 min of the recordings was similar between slices from translin KO mice and wildtype littermates (wildtype littermates: 150.03 ± 7.8%, n = 5; translin KO mice: 151.2 ± 13.4%, n = 5, t-test, *p* = 0.936). Furthermore, slices from translin KO mice showed no impairment in the FSK-LTP compared to the WT mice (**Fig. 2*B***; n = 5 for translin KO mice, n = 6 for wildtype littermates, two-way repeated-measures ANOVA, F_(1,9)_ = 0.07, *p* = 0.79). The average of the initial fEPSP slope over the last 20 min of the recordings was comparable between slices from translin KO mice and wildtype littermates (wildtype littermates: 180 ± 14.3%, n = 6; translin KO mice: 180.4 ± 11.8%, n = 5, t-test, *p* = 0.982).

**Figure 2.**
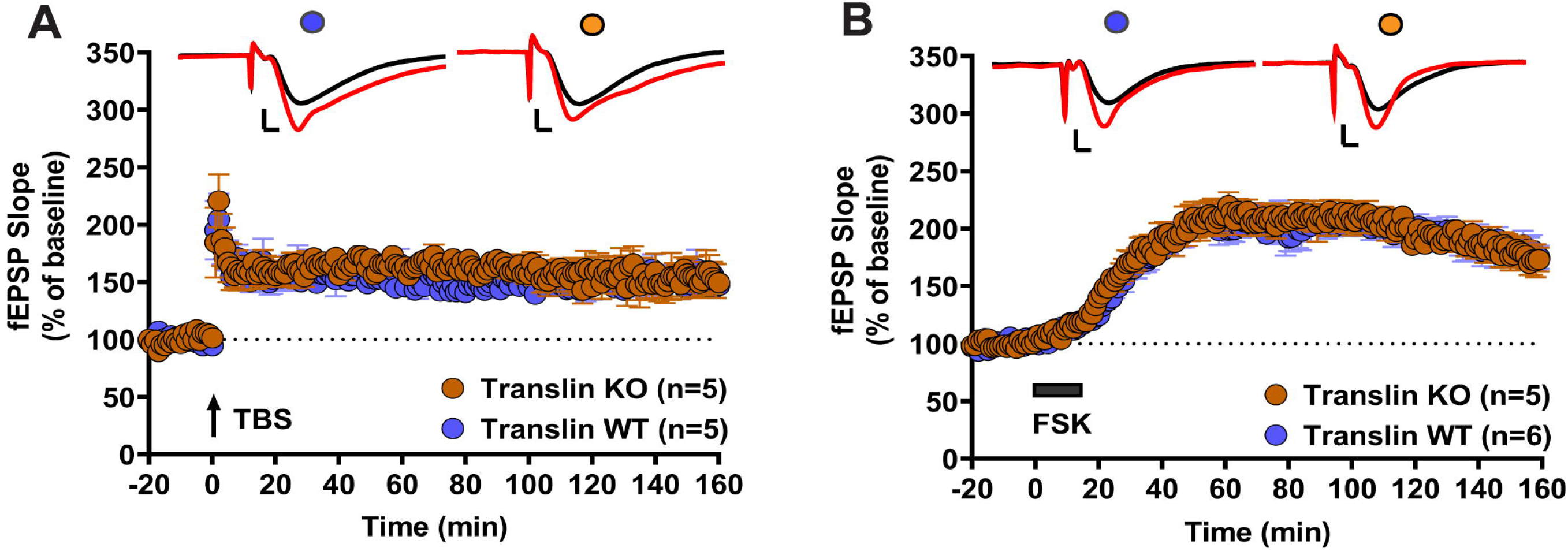
Translin KO mice show no alterations in predominantly presynaptic forms of long-lasting LTP. ***A***. Theta-burst stimulation (15 bursts of four 100 Hz pulses delivered at 5 Hz for a total of 3 s) induced similar levels of long-lasting LTP in slices from translin KO mice (n=5; 1 male, 4 females) or wildtype littermates (n=5; 2 males, 3 females) (two-way repeated measures ANOVA, F_(1,8)_ = 0.007, *p* = 0.94). ***B***. Slices from translin KO mice (n=5; all females) and wildtype littermates (n=6; 1 male, 5 females) displayed similar levels of forskolin (FSK)-induced long-lasting potentiation (two-way repeated-measures ANOVA, F_(1,9)_ = 0.07, *p* = 0.79). Representative traces before (black) and after (red) stimulation are shown on top of each graph. Scale bars for traces 2 mV, 2 ms. ‘n’ refers to the number of mice used. Error bars reflect S.E.M.

Taken together, these data suggest that translin is selectively involved in mediating the long-lasting form of LTP induced by spaced tetanic stimuli, but not in LTP induced by massed stimuli, TBS or forskolin.

### Translin KO mice exhibit unaltered mGluR-LTD and protein levels of hippocampal FMRP

One of the most well-studied RBPs is fragile X mental retardation protein (FMRP). Exaggerated metabotropic glutamate receptor-mediated LTD (mGluR-LTD) is a well characterized phenotype of FMRP KO mice and has been proposed as an underlying mechanism of fragile X syndrome (34-36). Because both translin/trax and FMRP mediate local protein synthesis (15, 36), we tested mGluR-LTD in hippocampal slices from translin KO mice. In contrast to the findings from FMRP KO mice, mGluR-LTD was unaffected in slices from translin KO mice (**Fig. 3*A***; n = 5 for each group, two-way repeated measures ANOVA, F_(1,8)_ = 0.08, *p* = 0.79). The average of the initial fEPSP slope over the last 20 min of the recordings was comparable between slices from translin KO mice and wildtype littermates (wildtype littermates: 75.9 ± 3.2%, n = 5; translin KO mice: 78.9 ± 3.1%, n = 5, t-test, *p* = 0.473). We reasoned that if translin and FMRP are functionally independent, loss of translin/trax should not cause a compensatory increase in FMRP protein levels. Indeed, Western blot analyses showed no changes in the protein levels of hippocampal FMRP in translin KO mice relative to wildtype littermates (**Fig. 3*B***; translin KO mice: 102.5 ± 0.4%, n = 6; wildtype littermates: 100 ± 4.3%, n = 6, t-test, *p* = 0.36). Our data indicate that translin/trax and FMRP play distinct roles in hippocampal synaptic plasticity.

**Figure 3.**
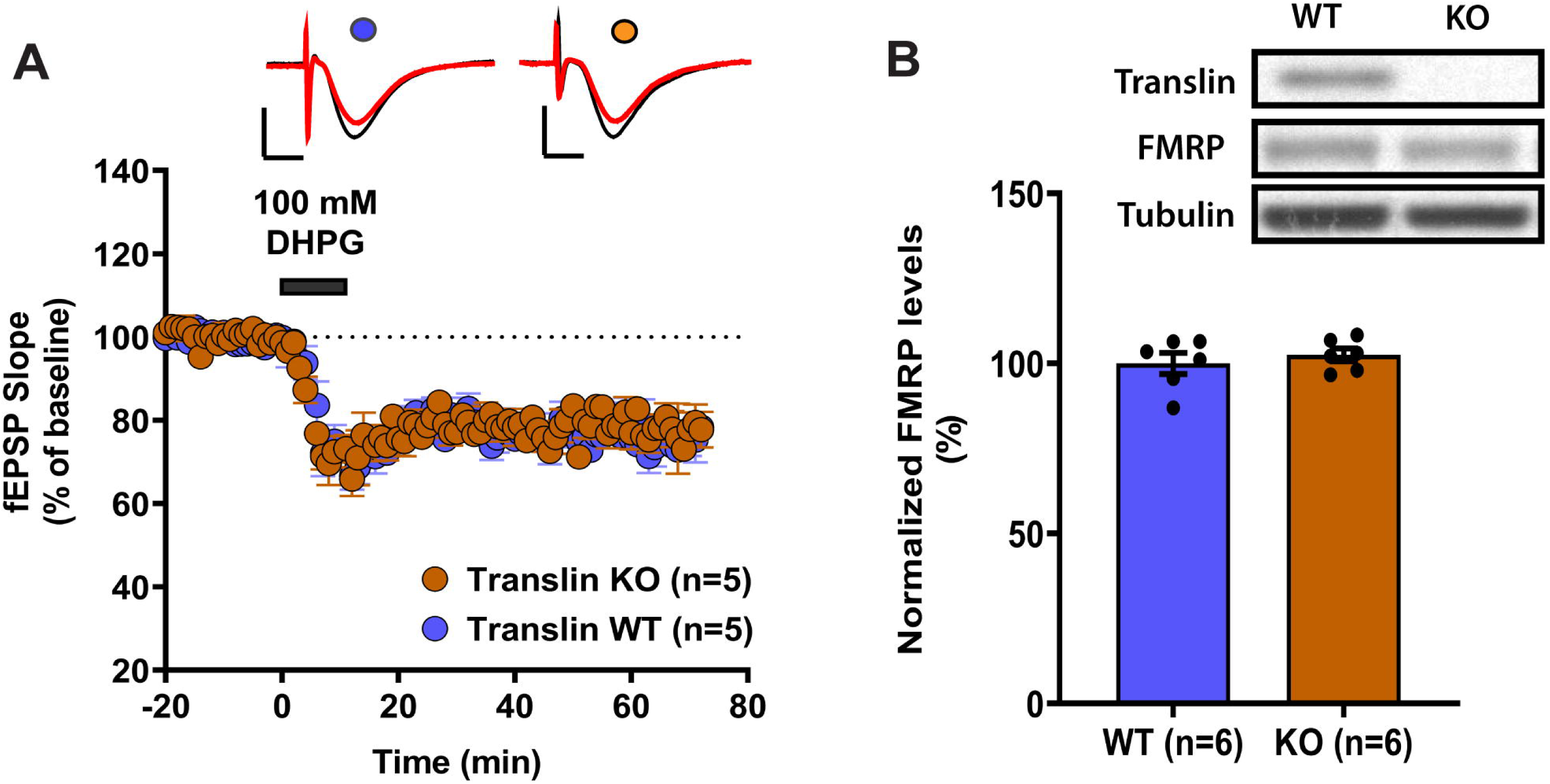
Translin KO mice show unaltered mGluR-LTD and unchanged hippocampal protein levels of FMRP. ***A***. Hippocampal slices from translin KO mice (n=5; males) and wildtype littermates (n=5; all males) displayed similar mGluR-LTD induced by bath application of 100 mM of DHPG for 10 minutes (two-way repeated measures ANOVA, F_(1,8)_ = 0.08, *p* = 0.79). Representative traces before (black) and after (red) stimulation are shown on top of the graph. Scale bars for traces 2 mV, 5 ms. ***B***. Hippocampal extracts from translin KO mice (n=6) and wildtype littermates (n=6) had similar protein levels of FMRP (t-test, *p* = 0.36). Beta-tubulin was used as the loading control and the expression level was normalized to the level of wildtype littermates. Representative blots are shown on top of the graph. ‘n’ refers to the number of mice used. Error bars reflect S.E.M.

## Discussion

The translin/trax complex is implicated in neuropsychiatric disorders (37). However, the role of translin/trax in synaptic plasticity is largely unknown. In a previous report (15), we provided the first evidence that translin/trax mediates activity-dependent synaptic translation that is critical for synaptic tagging and capture, a form of heterosynaptic associative plasticity (38, 39), and long-term memory. The present study further determined that translin/trax is selectively required for spaced tetani-induced LTP, a long-lasting form of hippocampal synaptic plasticity that is mediated by postsynaptic PKA activity.

We found that translin/trax is required for spaced 4-train-LTP that relies on postsynaptic PKA activity (28) but dispensable for TBS- and FSK-LTP that rely on presynaptic PKA activity (19, 28). These findings highlight the role of translin/trax in postsynaptic PKA signaling-dependent persistent synaptic plasticity. Notably, synaptic tagging and capture is impaired in the absence of translin/trax (15) but is intact when postsynaptically compartmentalized PKA signaling is disrupted (19). However, these studies used massed 4-train-LTP, which does not rely on PKA signaling, to induce synaptic tagging and capture. Considering this experimental caveat, multi-disciplinary approaches to manipulate localized translin/trax-PKA signaling with high spatiotemporal specification will further dissect the role of translin/trax in postsynaptic PKA signaling-mediated persistent synaptic plasticity.

We found that mice lacking translin/trax display electrophysiological phenotypes that are distinct from those observed in mice lacking FMRP, regardless of sharing some common behavioral abnormalities (22). Translin KO mice show deficits in spaced 4-train LTP (**Fig.1B**) and high-frequency stimulation-induced synaptic tagging and capture (15), but FMRP KO mice do not exhibit these impairments (36, 40). Exaggerated mGluR-LTD is the prominent phenotype of FMRP KO mice (35), but it was not observed in translin KO mice (**Fig. 3A**). Thus, our study demonstrates a selective role for translin/trax in synaptic plasticity and provides a foundation for future studies defining signaling pathways that enable synaptic stimulation to trigger the activation of this microRNA-degrading enzyme.

Based on our published and current findings, we propose a working model in which translin/trax mediates persistent synaptic plasticity (**Fig.4**). During basal synaptic transmission **(Fig.4A)**, the translin/trax microRNA-degrading enzyme is localized within the processing bodies (P-bodies) with its RNase inactive. This is supported by our previous data showing colocalization of trax with the P-body marker GW182 in hippocampal primary neuron dendrites (15). Given that P-bodies also contain mRNAs translationally repressed by the microRNA-mediated silencing complex (miRISC) (41-, the translin/trax complex is well-positioned to degrade microRNAs. Based on our preliminary data (not shown) suggesting the interaction of translin and a PKA-anchoring protein in hippocampal tissue, it is possible that PKA-anchoring proteins might tether translin/trax complex and PKA within P-bodies in postsynaptic compartments. Following persistent plasticity-inducing stimuli or learning **(Fig.4B)**, P-bodies are translocated to dendritic spines (44), and localized pools of PKA and the translin/trax complex are activated. Active translin/trax RNase then degrades microRNAs, relieving transcripts from translational repression. This allows the production of key plasticity-related proteins required for persistent plasticity. We have previously identified activin receptor type IC (ACVR1C) as one such plasticity-related protein (15). Whether PKA directly activates the translin/trax complex or acts through other targets warrants further investigation.

**Figure 4.**
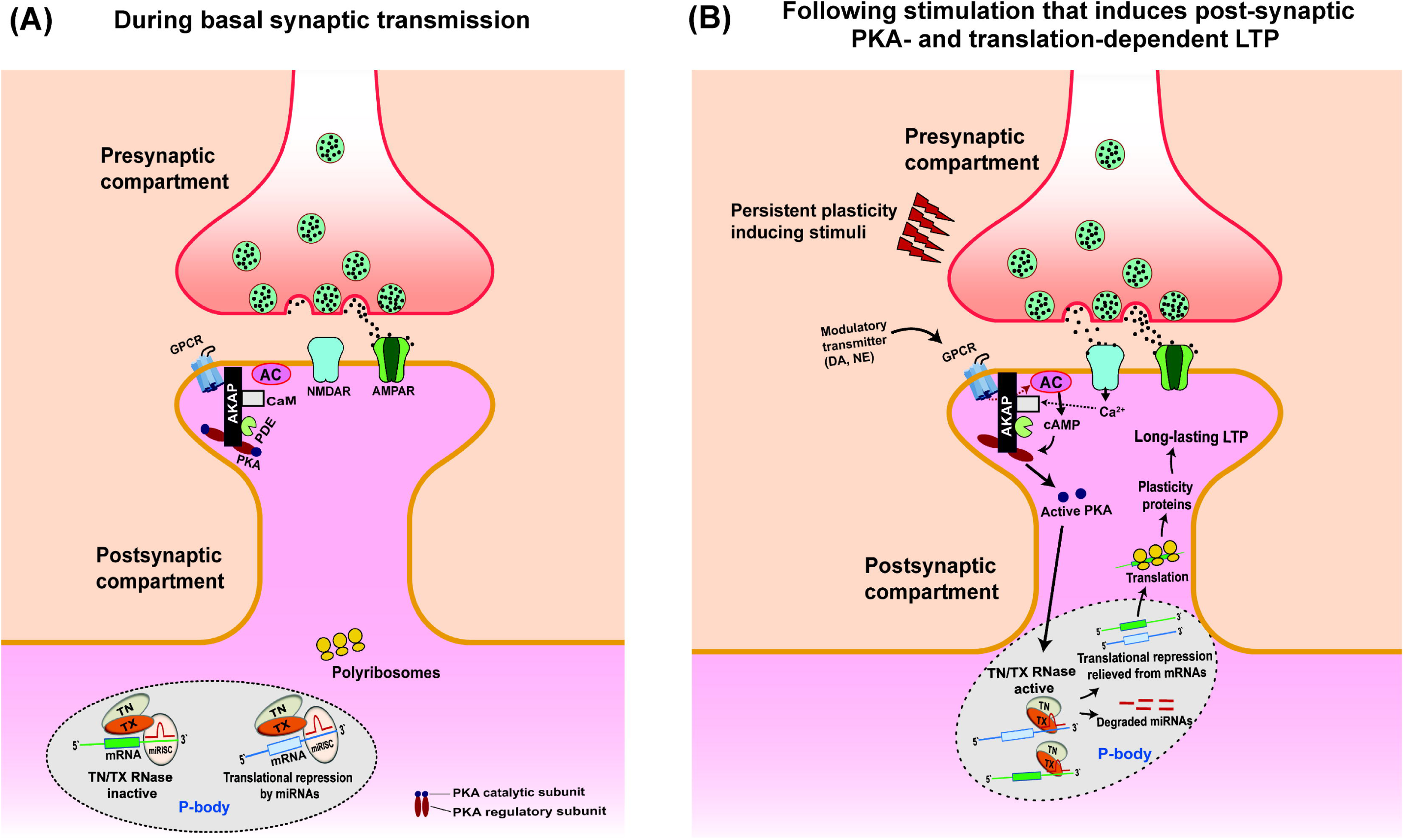
A working model for the role of translin/trax RNase complex in hippocampal synaptic plasticity. **(A)** During basal synaptic transmission, the inactive translin/trax (TN/TX) RNase complex localizes to the discrete ribonucleoprotein foci called Processing body (P-body) in the postsynaptic dendrites. P-bodies also contain the microRNA-induced silencing complex (miRISC) formed by the microRNAs and other associated protein factors that bind to the mRNA transcripts and repress their translation. **(B)** Synaptic activity that induces persistent plasticity leads to NMDAR-mediated Ca^2+^ entry and GPCR activation by modulatory neurotransmitters (e.g. norepinephrine and dopamine). This results in adenylyl cyclase activation, rise in cAMP levels and PKA activation in the postsynaptic compartment. Synaptic activity also leads to dynamic changes in P-bodies causing them to localize in the vicinity of active dendritic spines. Active PKA subsequently results in activation of the translin/trax RNase in the P-bodies, either directly or through other targets. Once active, the translin/trax RNase degrades microRNAs bound to the mRNA transcripts, thus reversing the translational silencing. The released mRNAs are then translated by the polyribosomes leading to the synthesis of key plasticity-related proteins required for long-lasting LTP.

Taken together, these findings expand our understanding of the role of translin/trax in persistent synaptic plasticity. Future investigations are needed to directly validate the proposed model. First, the compartment-specific function of translin/trax needs to be further validated using hippocampal subregion-specific deletion of translin/trax using Cre-dependent viral strategies. Second, experiments are also needed to validate the dynamics of translin/trax localization in P-bodies following plasticity-inducing stimuli. Tagging a fluorescent reporter to translin or trax would enable tracking the localization of translin/trax but achieving this without affecting molecular interactions and RNase activity is challenging. Lastly, molecular assays to determine mechanisms by which translin/trax interacts with localized PKA signaling will be required. Given the role of translin/trax in synaptic memory mechanisms, our findings pave the way for future research aimed at elucidating the pathophysiology of neuropsychiatric disorders.

## List of abbreviations

aCSF: artificial cerebrospinal fluid
AC: adenylyl cyclase
ACVR1C: activin receptor type 1-C
AMPAR: α-amino-3-hydroxy-5-methyl-4-isoxazolepropionic acid receptor
CaM: calmodulin
cAMP: cyclic adenosine monophosphate
DHPG: (*RS*)-3,5-Dihydroxyphenylglycine
fEPSP: field excitatory postsynaptic potential
FMRP: fragile X mental retardation protein
FSK: forskolin
GPCR: G-protein coupled receptor
LTP: long-term potentiation
LTD: long-term depression
mGluR: metabotropic glutamate receptor
miRISC: microRNA-induced silencing complex
NMDAR: N-methyl D-aspartate receptor
PDE: phosphodiesterase
PKA: protein kinase A
RBP: RNA-binding protein
RNase: ribonuclease
TBS: theta-burst stimulation
TN/TX: translin/trax
Trax: translin-associated protein-X

## Declarations

### 1. Ethics approval

All the experiments presented here were performed using mice. All experiments were performed according to the National Institutes of Health guidelines and were fully approved by the Institutional Animal Care and Use Committee at the University of Pennsylvania.

### 2. Consent for publication

Not applicable

### 3. Availability of data and materials

All the data supporting the conclusions of this article are included within the manuscript. Original data files and any supporting materials are available upon request, from the corresponding authors.

### 6. Authors’ contributions

A.J.P and T.A designed the experiments. A.J.P performed the electrophysiology and western blotting experiments. A.J.P performed data analysis. A.J.P and M.S.S prepared figures and wrote the manuscript with inputs from J.M.B and T.A. All the listed authors have read and approved the manuscript for publication.

## 7. Acknowledgements

Not applicable

